# An epilepsy-associated CILK1 variant compromises KATNIP regulation and impairs primary cilia and Hedgehog signaling

**DOI:** 10.1101/2024.05.14.594243

**Authors:** Ana Limerick, Ellie A. McCabe, Jacob S. Turner, Kevin W. Kuang, David L. Brautigan, Yi Hao, Cherry Chu, Sean H. Fu, Sean Ahmadi, Wenhao Xu, Zheng Fu

## Abstract

Mutations in human *CILK1* (ciliogenesis associated kinase 1) are linked to ciliopathies and epilepsy. Homozygous point and nonsense mutations that extinguish kinase activity impair primary cilia function, whereas mutations outside the kinase domain are not well understood. Here, we produced a knock-in mouse equivalent of the human *CILK1* A615T variant identified in juvenile myoclonic epilepsy (JME). This residue is in the C-terminal region of CILK1 separate from the kinase domain. Mouse embryo fibroblasts (MEF) with either heterozygous or homozygous A612T mutant alleles exhibited a higher ciliation rate, shorter individual cilia and up-regulation of ciliary Hedgehog signaling. Thus, a single A612T mutant allele was sufficient to impair primary cilia and ciliary signaling in MEFs. Gene expression profiles of wild type versus mutant MEFs revealed profound changes in cilia-related molecular functions and biological processes. CILK1 A615T mutant protein was not increased to the same level as the wild type protein when co-expressed with scaffold protein KATNIP (katanin-interacting protein). Our data show that KATNIP regulation of a JME-associated single residue variant of CILK1 is compromised and this impairs the maintenance of primary cilia and Hedgehog signaling.

## Introduction

Most eukaryotic cells use a microtubule-based apical membrane protrusion called the primary cilium to sense environmental cues and transduce extracellular signals [1]. The primary cilium functions as a signaling hub, compartmentalizing ion channels, receptor tyrosine kinases, G-protein coupled receptors, and second messengers (e.g. calcium and cyclic AMP) [2]. Primary cilia defects impair the sensory and signaling functions of a cell, leading to abnormalities in tissue development and homeostasis that underlie a large category of human disorders referred to as ciliopathies [3-5]. Patients with ciliopathies exhibit tissue and skeletal deformities, cognitive deficits and behavioral anomalies, implicating an important role of primary cilia as a non-synaptic mechanism to regulate neuronal circuits and functions [6].

One of the key regulators that control primary cilia formation and elongation is CILK1 (ciliogenesis associated kinase 1) [7]. CILK1 consists of an N-terminal catalytic domain (residues 1-284) and a C-terminal domain with an intrinsically disordered region (IDR, residues 285-632). The scaffold protein KATNIP binds to the IDR and stabilizes and activates CILK1 [8]. The homozygous mutations (e.g. R272Q) in the catalytic domain of CILK1 abolished kinase activity, impaired cilia length control and the Hedgehog signaling, and caused ciliopathy phenotypes [9]. The heterozygous variants in the IDR (e.g. K305T and A615T) are strongly linked to juvenile myoclonic epilepsy (JME) [10]. However, it remains to be determined whether these JME-associated heterozygous variants can exert a significant impact on primary cilia and by what mechanisms they would affect CILK1 functions.

## Results

### 1. Mutant cells with the mouse equivalent of human *CILK1* A615T

To evaluate the impact of an epilepsy-associated mutation *CILK1* A615T on primary cilia and ciliary Hedgehog signaling, we used CRISPR/Cas9 to genetically engineer mutations in exon 13 of mouse *Cilk1* (Fig. 1A). These knock-in mutations converted Ala612 (the mouse equivalent of human Ala615) to Thr and also introduced an Hpy99I restriction enzyme recognition site near the A612T mutation, as confirmed by sequencing results (Fig. 1B). Restriction digestion of PCR products by Hpy99I allowed us to distinguish between *Cilk1* wild type (WT) and A612T mutant alleles, as shown in genotyping results (Fig. 1C). We subsequently isolated mouse embryonic fibroblasts (MEFs) from E15.5 *Cilk1* WT and mutant embryos with heterozygous and homozygous A612T substitutions.

**Figure 1:**
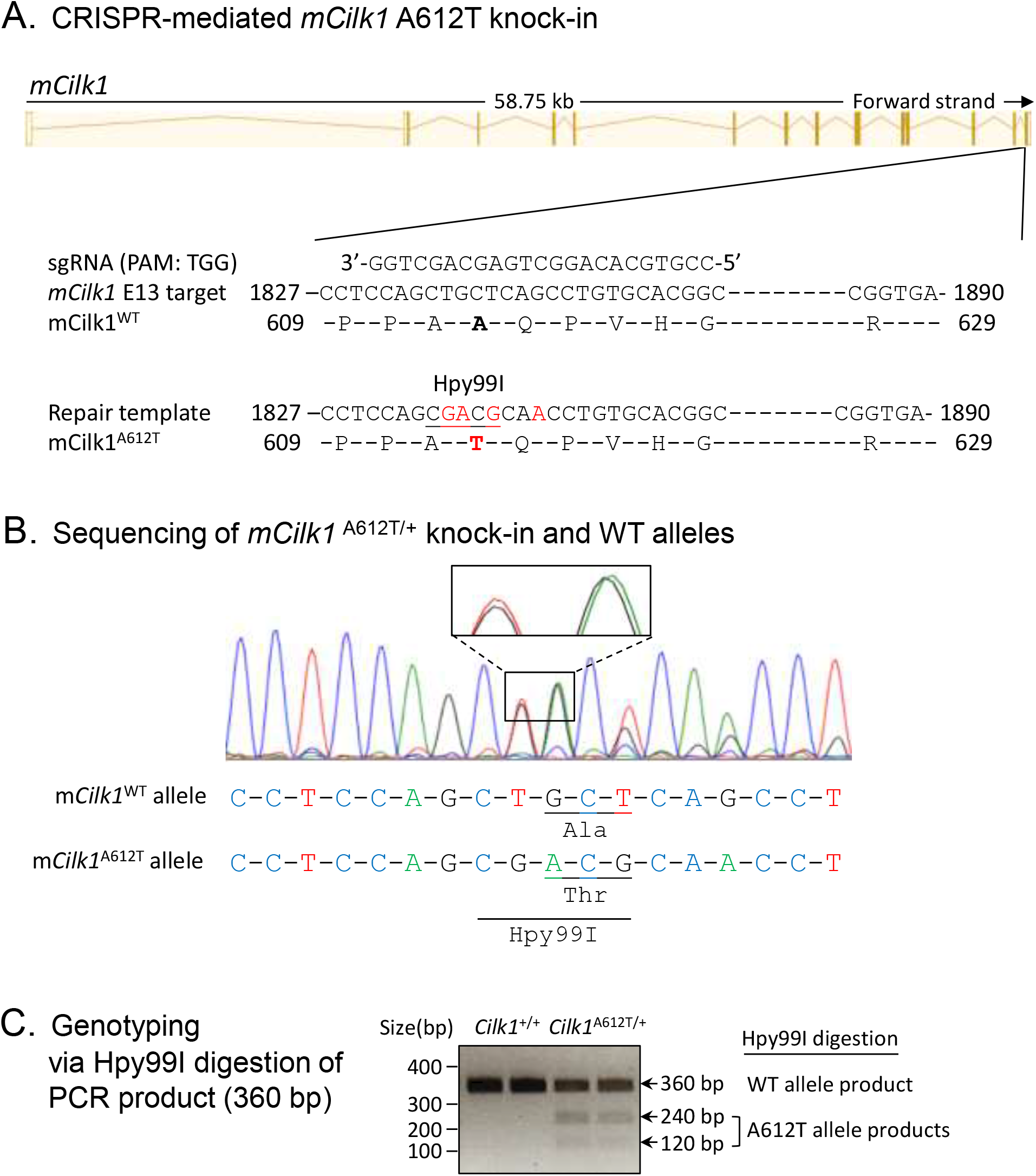
Generation of the *Cilk1* A612T knock-in mouse model. **(A)** A schematic illustration of CRISPR/Cas9-mediated knock-in of the A612T mutation and an Hpy99I restriction enzyme cutting site into the *mCilk1* mutant allele. **(B)** Sequencing results confirming designed mutations that not only converted Ala612 to Thr but also introduce a new Hpy99I site near A612T for genotyping. One silent base substitutions (CAG→CAA) was also engineered to deter the recut of the repair template by Cas9 nuclease. **(C)** Genotyping results from Hpy99I-digested PCR products that can distinguish between the WT allele and the A612T mutant allele.

### 2. *Cilk1* A612T mutant allele induces higher ciliation frequency but shorter cilia

We immunostained MEFs with primary cilia marker Arl13B and basal body marker γ-tubulin (Fig. 2A), calculated percent of ciliated cells, and measured cilia length. Our results revealed that about 20% of WT MEFs had cilia under standard growth conditions (Fig. 2B). By comparison, MEFs carrying a heterozygous A612T mutation showed a statistically significant increase in the fraction of cells with cilia (ciliation rate) to nearly double the rate of WT MEFs (Fig. 2B). With both copies of the CILK1 gene mutated in homozygous MEFs there was a further increase in the fraction of cells with cilia, more than double compared to WT MEFs (Fig. 2B). These results indicated a dose-dependent effect of the amount of functional CILK1 required to control the formation of primary cilia. On the other hand, the average cilia length was 2.2 µm in WT MEFs compared to 1.8 µm in A612T mutant MEFs, a statistically significant decrease of about 18% (Fig. 2C). In this case MEFs bearing heterozygous A612T mutation were not significantly different from those bearing homozygous A612T mutation. This demonstrates haploinsufficiency of the WT allele, indicating that the length of the primary cilium requires both WT copies of the gene.

**Figure 2:**
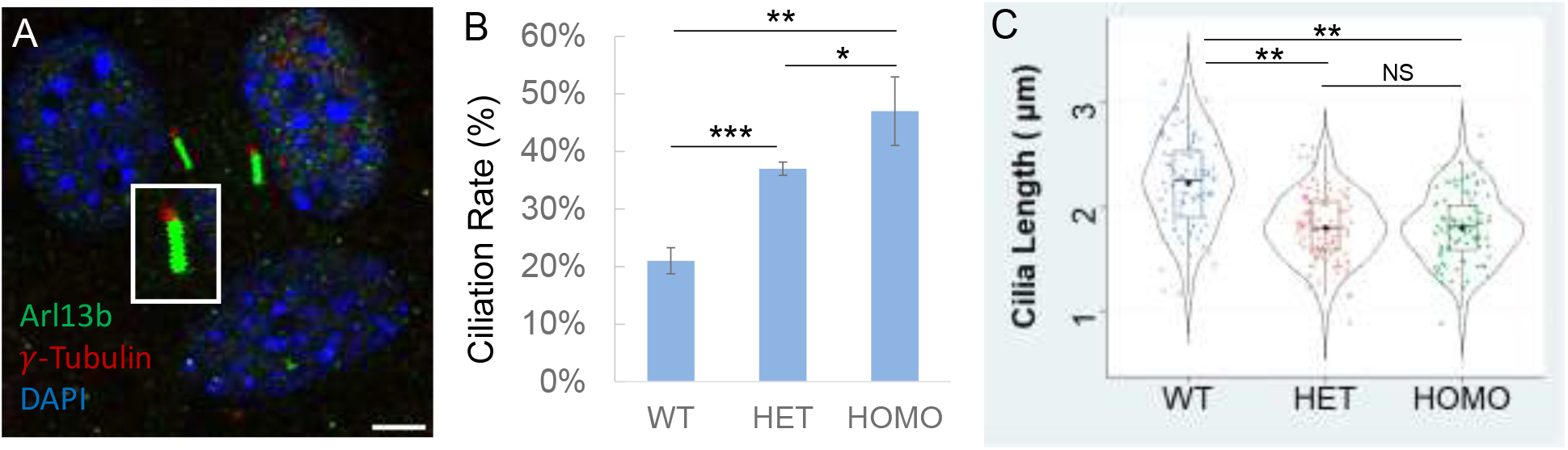
Effects of Cilk1 A612T mutant on primary cilia frequency and length. **(A)** Cilk1 WT and A612T mutant MEFs (mouse embryonic fibroblasts) were immunolabelled with the Arl13B antibody for primary cilia and the γ-Tubulin antibody for basal bodies, and stained with DAPI for nuclei. Scale bar, 3µm. **(B)** Shown are ciliation rates in MEFs of wild type (WT, 1167 cells), A612T heterozygous (HET, 1295 cells), and homozygous (HOMO, 1428 cells), mean ± SD, n = 3 independent experiments, Student’s *t*-test, *P < 0.05, **P < 0.01, ***P < 0.001. **(C)** A violin-box plot showing the distribution of numerical values of cilia length in Cilk1 WT control (n = 93 cilia), A612T HET (n = 109 cilia), and A612T HOMO (n = 94 cilia) MEFs, median with interquartile range box and min/max whiskers. One-way ANOVA followed by post-hoc Tukey HSD test was used to assess the significance of differences between pairs of group means. The significance level was at p values of 0.05 (*) and 0.01 (**). NS = not significant at both P values.

### 3. *Cilk1* A612T mutant allele triggers upregulation of the Hedgehog signaling

One test of cilia function is Hedgehog (Hh) signaling. The binding of an Hh ligand to its receptor Patched 1(Ptch1) relieves Ptch1-mediated inhibition of Smoothened (Smo), causing Smo to translocate to the primary cilium for activation of downstream effectors such as Gli1 and Gli2.

Smo translocation and activation is also induced by Hh agonists such as SAG. We treated wild type MEF cells with either DMSO as control or SAG (50 nM and 200 nM) to activate Hh signaling (Fig. S1). We observed a significant accumulation of Smo in the primary cilium upon treatment with 200 nM SAG, based on Smo co-localization with the cilia marker Arl13B (Fig. 3A and Fig. S1). In comparison to WT cells, both heterozygous and homozygous A612T mutant cells showed greater up-regulation of Smo upon SAG stimulation. We confirmed there was no major difference in the protein level of Smo and Gli1 between the heterozygous and the homozygous mutant cells. These results indicated that one *Cilk1* A612T mutant allele was sufficient to up-regulate the Hedgehog pathway.

**Figure 3:**
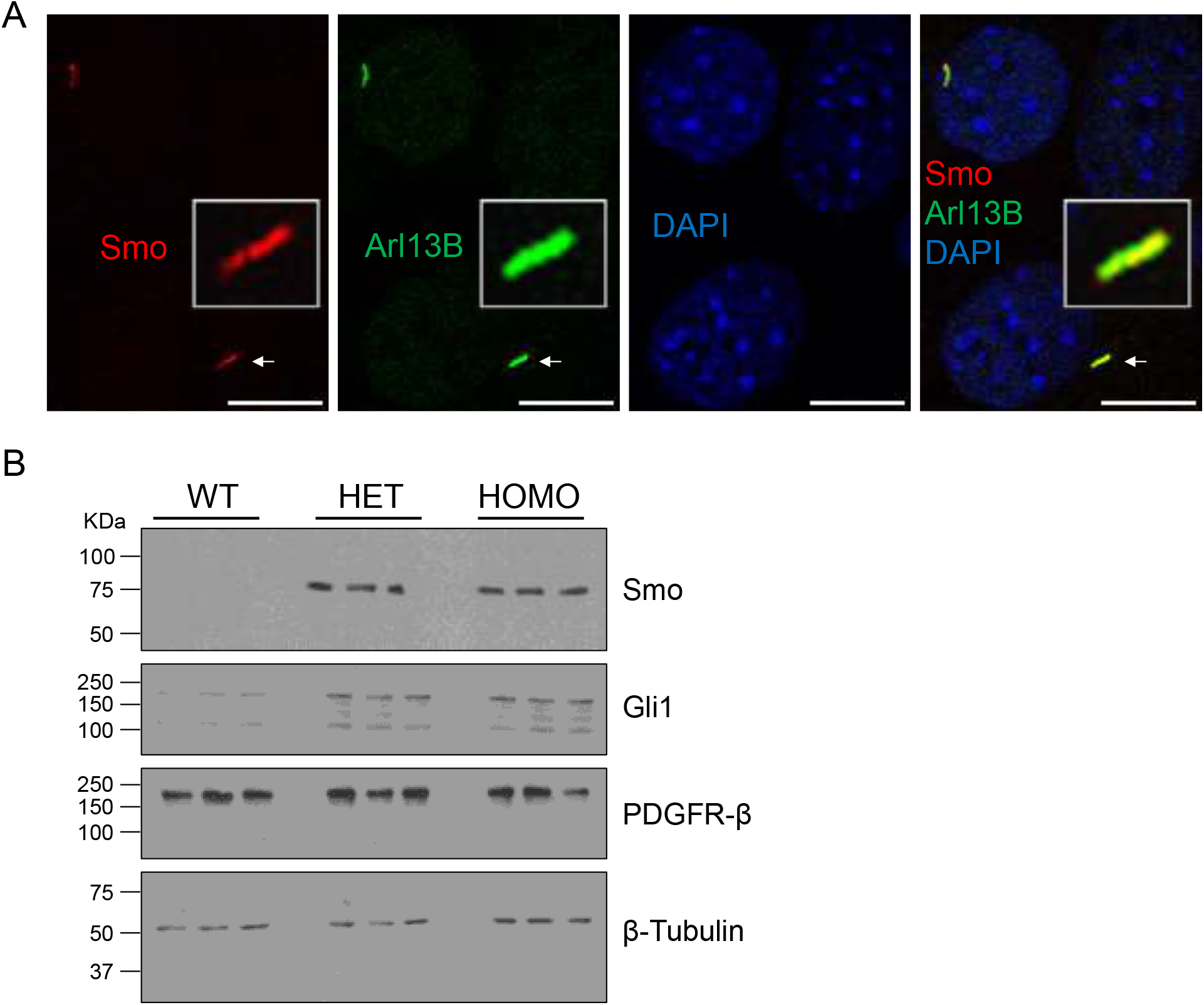
Effects of Cilk1 A612T on the Hedgehog signaling. **(A)** MEFs were treated with Smo agonist SAG (200 nM) to activate the Hedgehog signaling. Shown are MEFs immunolabelled with Smo and Arl13B antibodies and the overlay of Smo and Arl13B signals in primary cilia. Scale bar, 3 µm. **(B)** Equal amount of total proteins extracted from MEFs treated with SAG were blotted with Smo, Gli1, PDGFRβ, and β-Tubulin (the loading control) antibodies. WT = Cilk1 wild type; HET = Cilk1 A612T heterozygous; HOMO = Cilk1 A612T homozygous.

### 4. Cilk1 A162T variant produced profound alterations in cilia-related receptor tyrosine kinase (RTK) pathways and biological functions

To further examine the effects on cellular functions and signaling pathways, we applied RNA-seq analysis to CILK1 WT MEFs and A612T mutant MEFs and calculated the fold change of gene expression in mutant cells relative to WT cells. A volcano plot of differentially expressed genes (DEG) showed that 1286 DEG were identified including 931 up-regulated genes and 355 down-regulated genes (Fig. 4A, supplementary table 1). We performed Gene Ontology (GO) enrichment analysis of DEG (Fold Change >=2 and FDR < 0.001). Molecular function (MF) analysis revealed enrichment of gene expression changes in transmembrane receptor kinase activity (Fig. 4B), which is consistent with role of primary cilia in the compartmentalization and regulation of receptor tyrosine kinase pathways. Biological Process (BP) analysis identified that these changes are mainly involved in the regulation of extracellular matrix organization, angiogenesis, and cell migration (Fig. 4C). Importantly, both MF and BP analyses suggested that these changes are related to transmembrane receptor tyrosine kinase pathways and protein phosphorylation.

**Figure 4:**
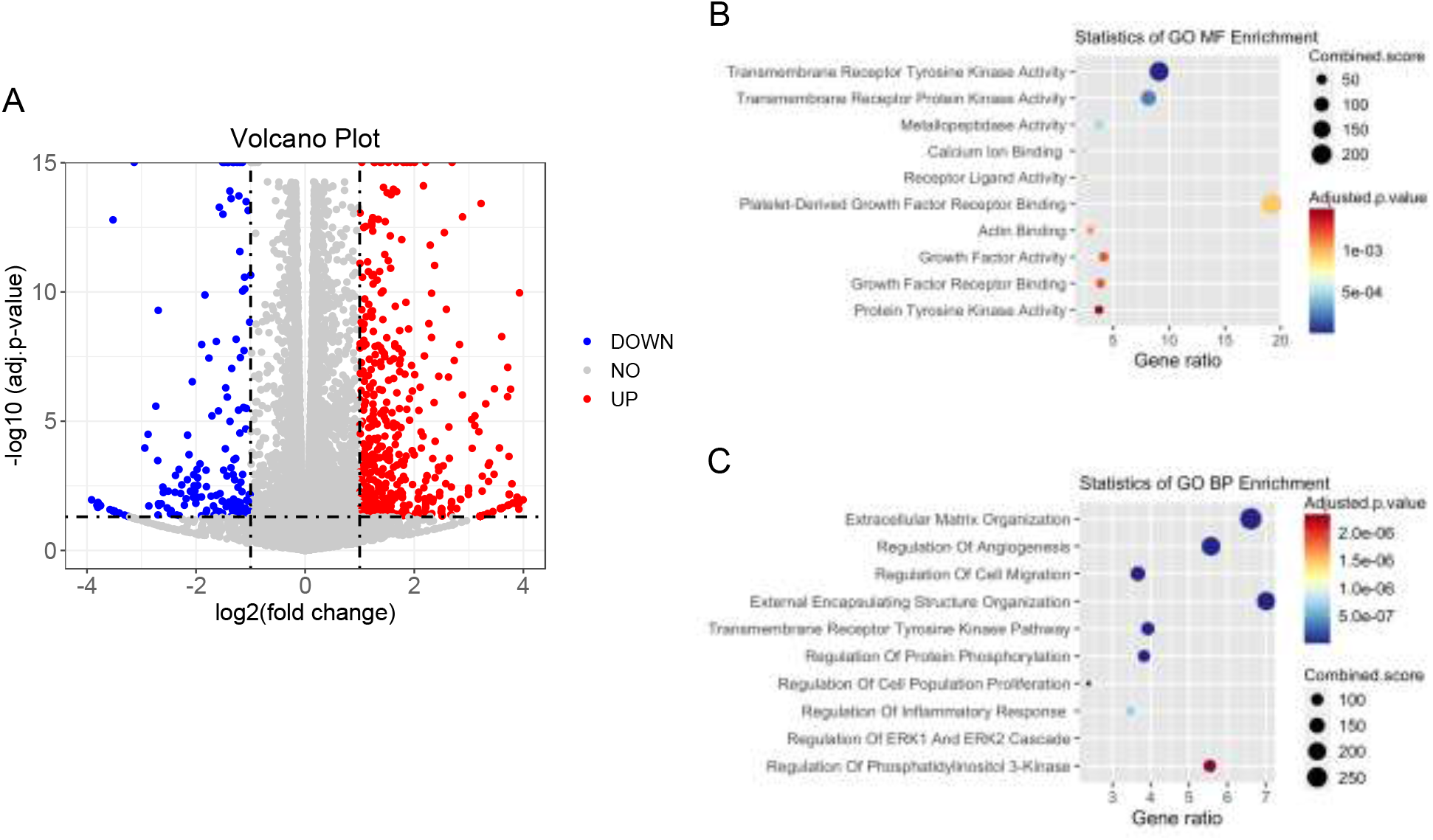
RNA-seq analysis of Cilk1 WT and A612T mutant MEFs. Total RNA was extracted and purified from MEF cells expressing wild type (WT) or homozygous A612T mutant Cilk1. (A) The volcano plot showing differentially expressed genes (fold change >2 or <0.5 and p-value < 0.05) in A612T mutant cells compared to WT cells; (B) The molecular function (MF) analysis of differentially expressed genes; (C) The biological Process (BP) analysis of differentially expressed genes.

### 5. The A615T variant compromised KATNIP regulation of CILK1 protein

Recently, we showed that the scaffold protein KATNIP interacts with CILK1, increasing protein levels and activation of the kinase [8]. Given that the A615T mutation is located in the intrinsically disordered region of CILK1 which mediates CILK1 interaction with KATNIP, we examined if the A615T mutation impairs KATNIP regulation of CILK1. We transfected HEK293T cells with Flag-CILK1 (WT or A615T) and co-transfected an increasing amount of Flag-KATNIP, and then quantified the protein levels of CILK1 on Western blots. While KATNIP enhanced WT CILK1 levels over 20-fold in a dose-dependent manner, at the same doses it was 2-fold less effective at increasing the protein level of A615T mutant protein (Fig. 5). This result indicates that the A615T variant was deficient in the KATNIP regulation of CILK1 protein, consistent with the lack of control of ciliogenesis by this mutant protein (Fig. 2).

**Figure 5:**
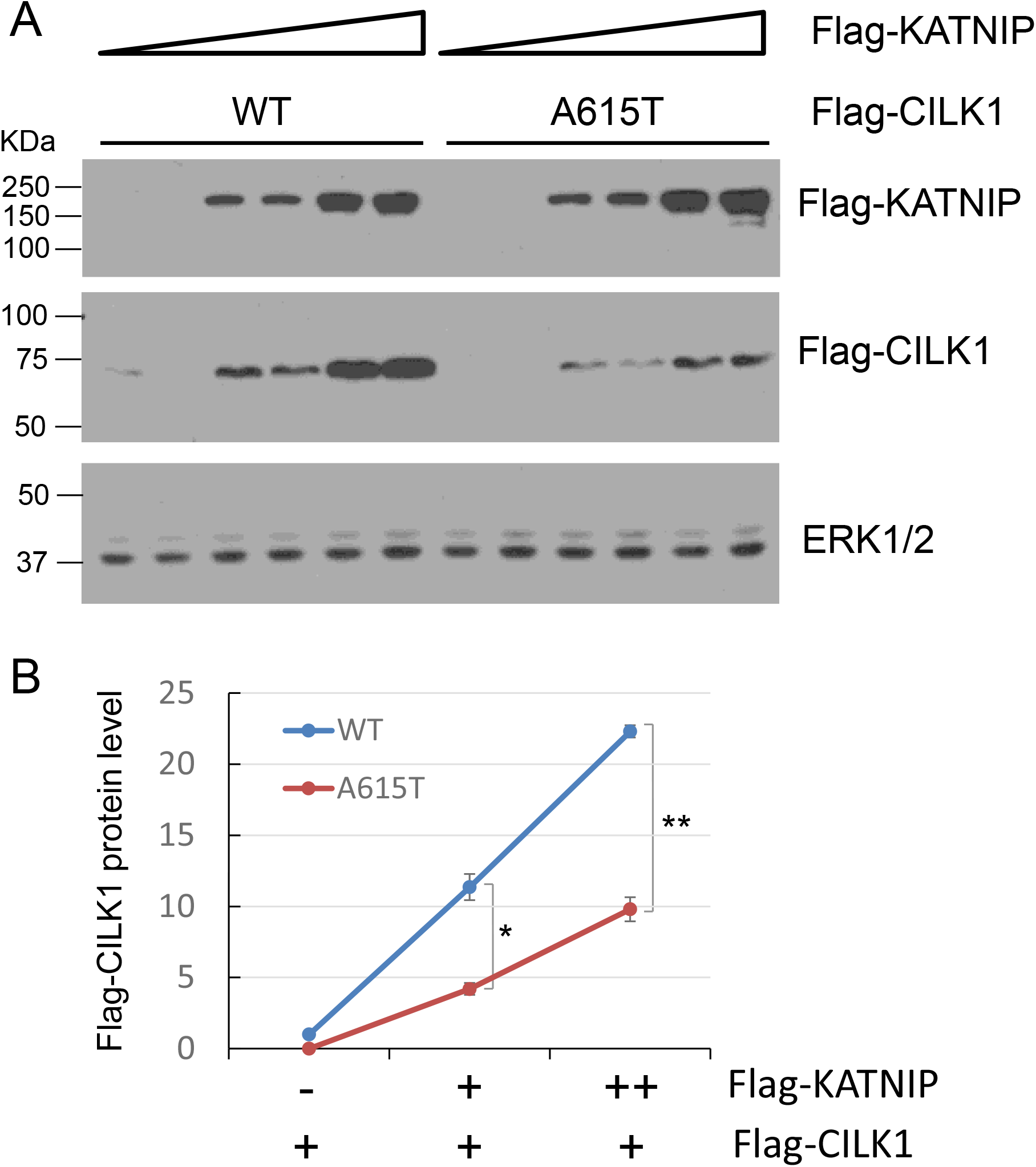
Effect of CILK1 A615T mutation on CILK1 stabilization by KATNIP. **(A)** Flag-CILK1 WT or A615T mutant was co-expressed with an increasing amount of Flag-KATNIP in HEK293T cells. Equal amount of total proteins from cell lysates were blotted with Flag and ERK1/2 antibodies. **(B)** Flag-CILK1 proteins normalized against total ERK proteins were plotted as a function of the increasing level of Flag-KATNIP proteins. Quantification data were shown as mean ± SD, two-tailed Student t-test, *p < 0.05, **p < 0.01. Shown here is the representative data from three independent experiments.

## Discussion

We examined the effects of a knock-in single residue substitution A612T in the non-catalytic region of CILK1 kinase on its function in MEFs. Our results showed significantly shorter cilia in both heterozygous and homozygous A612T mutant MEFs that was significantly different from wild type MEFs (Fig. 2). Previous studies of CILK1 null mice revealed that when one copy of the *Cilk1* gene was deleted, the remaining copy was sufficient to support a wild-type phenotype in MEFs. Thus, CILK1 shows haplosufficiency and ciliopathy phenotypes in MEFs require deletion or inactivation of both *Cilk1* alleles [9, 11, 12]. In contrast, heterozygous *Cilk1A612T*^+/-^ MEFs exhibited shorter cilia and increased ciliation compared to WT MEFs, so we propose this was due to a dominant negative effect of the mutant allele. Somehow the mutant protein interferes with the function of the WT protein in the same cells to produce a cilia phenotype. One means of producing a dominant negative effect is to compete for a limiting regulatory partner. In the current case this might be the activating subunit KATNIP.

Our data showed that when co-expressed with KATNIP the CILK1 A615T mutant protein levels were significantly lower compared to the wild type protein. This suggests the A615T mutation prevented KATNIP stabilization of CILK1 (Fig. 5). This could be due to weaker or non-productive binding of KATNIP to CILK1 A612T. If weaker, then there would not be dominant negative effects, however if KATNIP binding to CILK1 A612T was the same or even higher affinity than to WT then the mutant protein would deplete the KATNIP available for activating CILK1. These alternative mechanisms merit further investigation. Of course, it is also possible that the A615T mutant protein may gain functions yet to be identified to interfere with the control of cilia formation.

Our data clearly demonstrated strong effects of the CILK1 A615T variant on primary cilia and cilia-dependent Hedgehog signaling in MEFs, however its impact on neuronal functions and seizures and the underlying mechanisms in vivo remain unknown. The A612T mice are viable and fertile, so we will have a cohort of animals to study the neuronal and behavioral phenotypes. It is worth pointing out that alterations in primary cilia have been associated with various neurological disorders [6, 13, 14]. For instance, diminished primary cilia formation has been observed in psychiatric diseases such as schizophrenia and bipolar disorders [15]. Both spontaneous and induced seizures have been shown to disrupt primary cilia growth and length control [16]. Therefore, the primary cilium has been proposed as a possible marker for diagnosis and has the potential use for high-throughput drug screen in neurological diseases.

## Materials and Methods

### Reagents and Antibodies

pCMV6-Myc-Flag-CILK1 (RC213609) and pCMV6-Myc-Flag-KATNIP (RC220412) were from Origene (Rockville, MD, USA). CILK1 A615T mutant sequence was amplified by PCR and cloned into pCMV6-Myc-Flag vector at Custom DNA Constructs LLC (Mount Sinai, NY, USA). Smo (E-5) mouse monoclonal antibody (sc-166685) and Gli-1 (C-1) mouse monoclonal antibody (sc-515751) were from Santa Cruz Biotechnology (Dallas, TX, USA). PDGFRβ (28E1) rabbit monoclonal (#3169) and beta-Actin (13E5) rabbit monoclonal (#4970) antibodies were from Cell Signaling Technology (Danvers, MA, USA). Arl13B rabbit polyclonal antibody (17711-1-AP) and γ-tubulin mouse monoclonal antibody (66320-1-Ig) were from Proteintech (Rosemont, IL, USA). Goat anti-rabbit IgG (Alexa Fluor 488) antibody (ab150081) and goat anti-mouse IgG (Alexa Fluor 594) antibody (ab150120) were from Abcam (Cambridge, MA, USA). SAG (Smoothened Agonist) HCl (#S7779) was from Selleckchem Chemicals LLC (Houston, TX, USA).

### MEF cell isolation and culture

MEF cells were isolated from E15.5 embryos and maintained at 37°C and 5% CO2 in Dulbecco’s modified Eagle’s medium (DMEM) supplemented with 4.5 g/L glucose, 10% fetal bovine serum, and penicillin-streptomycin using a standard protocol (Durkin ME, 2013).

### Immunoblotting

Cells were lysed in lysis buffer (50 mM Tris-HCl, pH 7.4, 150 mM NaCl, 1% NP-40, 2 mM EGTA, complete protease inhibitors (Roche), 10 mM sodium orthovanadate, 5 mM sodium fluoride, 10 mM sodium pyrophosphate, 10 mM β–glycerophosphate, and 1 µM microcystin LR). Cell lysate was cleared by microcentrifugation. Cell extracts were boiled for 5 min in an equal volume of 2X Laemmli sample buffer and resolved by SDS-PAGE. Samples were transferred to a nitrocellulose membrane and blocked for one hour in 5% dry milk before primary antibody incubation in TBS containing 0.1% Tween-20 and 5% bovine serum albumin (BSA) for overnight at 4°C. This was followed by multiple rinses and one-hour incubation with horseradish peroxidase (HRP)-conjugated secondary antibody. Chemiluminescence signals were developed using Millipore Immobilon ECL reagents.

### Immunofluorescence

MEF cells grown on gelatin-coated coverslips were fixed by 4% paraformaldehyde in PBS, rinsed in PBS, and then permeabilized by 0.2% Triton X-100 in PBS. After one hour in blocking buffer (3% goat serum, 0.2% Triton X-100 in PBS), cells were incubated with primary antibodies at 4 °C overnight followed by rinses in PBS and one hour incubation with Alexa Fluor-conjugated secondary antibodies. After multiple rinses, slides were mounted in antifade reagent containing DAPI (4’,6-diamidino-2-phenylindole) for imaging via a confocal Laser Scanning Microscopy 700 from ZEISS (Chester, VA, USA) at the UVA Advanced Microscopy Facility.

### Cilia length measurement

The Zen 2009 program was used with a confocal Laser Scanning Microscope 700 from ZEISS (Chester, VA, USA) to collect z stacks at 0.5 μm intervals to incorporate the full axoneme based on immunostaining of cilia marker Arl13b and basal body marker γ-Tubulin. All cilia were then measured in ImageJ via a standardized method based on the Pythagorean Theorem in which cilia length was based on the equation L2 = z2 + c2, in which “c” is the longest flat length measured of the z slices and “z” is the number of z slices in which the measured cilia were present multiplied by the z stack interval (0.5 μm).

### RNA-seq analysis and enrichment analysis

RNA-seq analysis was conducted at the University of Virginia, Genome Analysis and Technology Core, RRID:SCR_018883. RNA-Seq reads were aligned using STAR (*Dobin et al. 2013*) with mouse genome build mm10 as reference. Expression count matrix were calculated using RSEM (*Li and Dewey 2011*). Differential expression analysis of RNA-seq was performed in R using the EdgeR package (*Robinson et al. 2010*) with a Benjamini–Hochberg FDR of 0.001. Three replicates were sequenced and analyzed for each condition of WT and A612T mutant. Ontology (BP and MF) and KEGG pathway enrichment analysis were performed by Enrichr (*Kuleshov et al. 2016*). Differentially-expressed genes with abs(log2FoldChane) >= 2 and FDR < 0.001 were used in the functional analysis. Volcano plot and enrichment plots were made in R (4.2.3).

### Statistical analysis

Experimental data were analyzed by the two-tailed Student’s t test to compare the means of two groups, and p values less than 0.05 were considered as significant. One-way ANOVA and post-hoc Tukey HSD test was used to assess the significance of differences between pairs of group means. For the Tukey test, the significance level at p values of 0.05 and 0.01 was used for evaluation.

## Acknowledgement

We thank the following core facilities at the University of Virginia School of Medicine for technical support: Genetically Engineered Murine Model, Advanced Microscopy Facility, and Genome Analysis and Technology. This work was supported by the National Institute of General Medical Sciences grant GM127690 to Z.F. and the National Cancer Institute Cancer Center Support Grant 5P30CA044579 to UVA Comprehensive Cancer Center Cores.

---

**Supplemental Figure 1:**
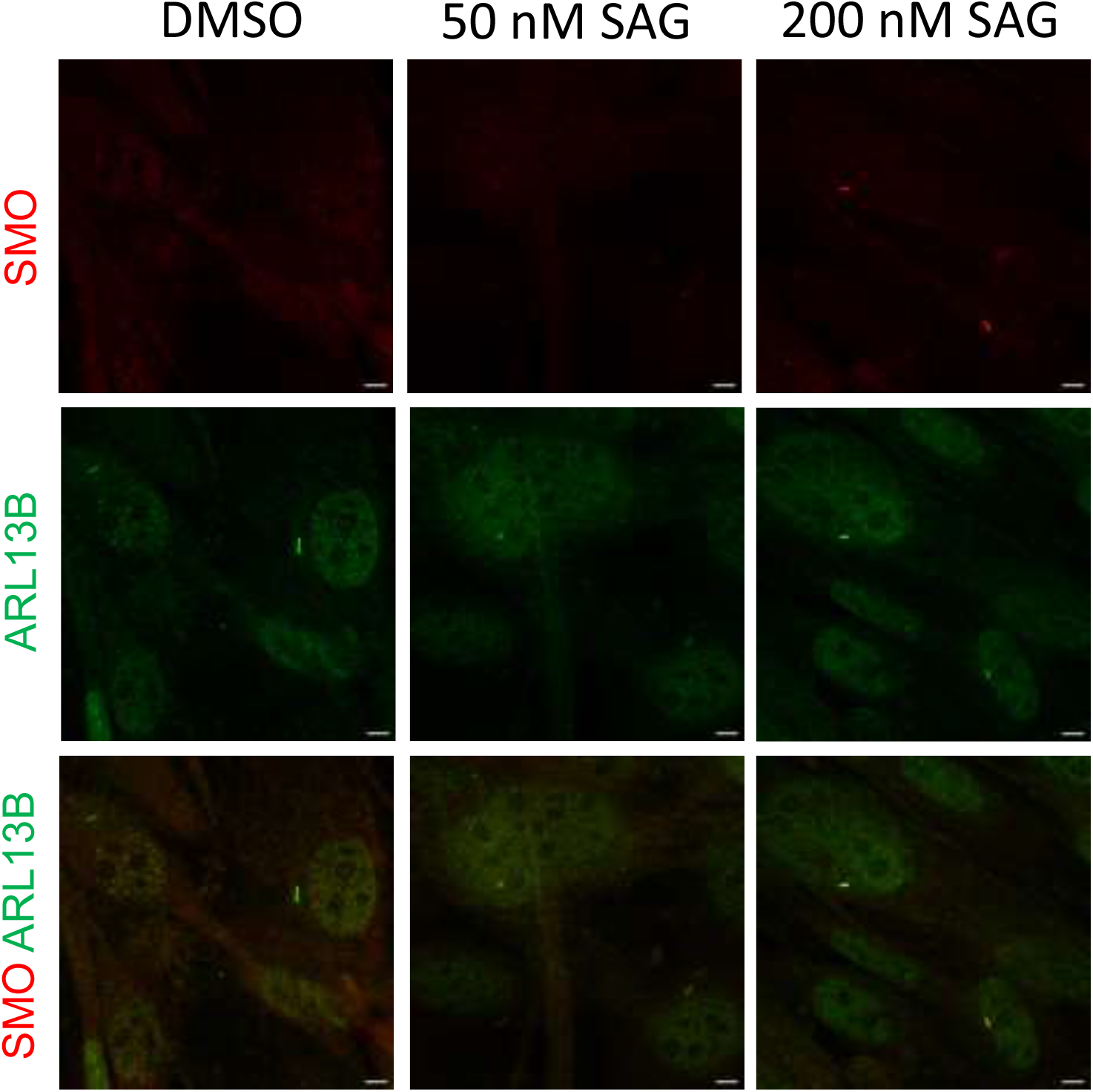
Smo ciliary localization induced by SAG. MEF cells were treated with either DMSO (control) or the Hh agonist SAG (50 nM, 200 nM). Cells were co-immunolabelled with antibodies against Smo and the cilia marker Arl13B to examine Smo localization in the primary cilium.

